# Ethological principles predict the neuropeptides co-opted to influence parenting

**DOI:** 10.1101/064808

**Authors:** Christopher B. Cunningham, Majors J. Badgett, Richard B. Meagher, Ron Orlando, Allen J. Moore

**Author notes:** Corresponding author: Allen J. Moore.

## Abstract

Ethologists predicted that parental care evolves by modifying suitable behavioural precursors in the asocial ancestor, such as nest building, defensive and aggressive behaviours, and potentially shared resources. From this, we predicted that the evolved mechanistic changes would reside in genetic pathways underlying these behavioural precursors. We tested this by measuring differential expression of neuropeptides in female *Nicrophorus vespilloides* Parenting in this species is extensive and complex as caring adults regurgitate food to begging, dependent offspring. We identified neuropeptides associated with mating, feeding, aggression, and social interactions by sampling females in different behavioural states: solitary, actively parenting, or post-parenting and solitary. We measured peptide abundance in adult female brains and identified 130 peptides belonging to 17 neuropeptides. Of these 17, seven were differentially expressed. Six of the seven were up-regulated during parenting. None of the identified neuropeptides have previously been associated with parental care, but all have known roles in the behavioural precursors. Two, tachykinin and sulfakinin, influence multiple pathways. Our study supports the prediction that appropriate behavioural precursors are likely targets of selection during the evolution of parenting. Evolutionary principles predicted neuropeptides influencing social behaviour, and our results provide several new candidate neuropeptides underpinning parenting.

---

The selective pressures that lead to the evolution of parental care are well documented. Parental care typically evolves to minimize unusually stressful or hazardous environments for offspring^1-3^. Although this hypothesis for the source of natural selection resulting in the evolution of parenting is widely supported^3^, parental care is not the only evolutionary solution to adverse conditions. Moreover, it may not be the most likely response as the evolution of parenting reflects changes in multiple behavioural inputs, involving many pathways^4^. At a minimum the evolutionary transition from asociality to subsociality involving direct parental care is predicted to require modification of the tendency to disperse from a mating site, a pause in reproduction and mating, defensive aggression to protect offspring and resources, changes in feeding behaviour, and a tolerance of increased social interactions^1-3,5^. Early ethological literature therefore predicts that parental care evolves only when there are suitable behavioural precursors present within the evolutionary ancestor, such as nest building, defensive postures and appropriately directed aggressive behaviours, and potentially shared resources^1,2^.

Despite these early predictions of the specific behaviours to be modified, the mechanistic alterations involved are relatively unknown. However, the predictions of ethologists imply expected underlying genetic pathways. In addition, Wright’s theory of nearly universal pleiotropy^6^, along with the ubiquity of regulatory evolutionary changes^7-9^, suggests that coopting behaviours will result in altered gene expression rather than the evolution of novel genes. Identifying the nature of selection can be useful for predicting the genetic changes underlying the evolution of social behaviour generally^5,10,11^. Therefore, we predict that parenting will involve changes in gene expression influencing feeding, mating, aggression, and increased tolerance for social interactions as these are the behaviours modified as lineages evolve from asocial to subsocial^1,2^.

Neuropeptides strongly influence the social behaviour of animals^12^ and many neuropeptides are likely to be associated with parenting. One of the most studied neuropeptides, oxytocin, is necessary for parenting across the animal kingdom^14^. There is a casual relationship between the neuropeptide *galinin* and parental care in mice^15^. We have recently provided evidence that at the transcriptional level *neuropeptide F receptor* is differentially expressed between parenting and non-parenting states in the burying beetle *Nicrophorus vespilloides*^16^. Moreover, individuals expressing parental care must undergo many rapid shifts in behaviour. Neuropeptides can exhibit their influence within minutes, have highly localized effects targeting very select neural circuits, or have highly widespread effects targeting many and diffuse neural circuits^16^. However, transcriptomics is not a particularly powerful method for identifying changes in neuropeptide expression. Neuropeptides generally have low gene expression^17^, highly restricted sites of release^16^, and can be hard to detect with transcriptomic studies that are not highly tissue specific^18^. Proteomics can overcome some of these limitations and provides a method to target proteins of interest.

Here, we test the hypothesis that a transition from a non-parenting state to a parenting state will reflect differences in expression of neuropeptides known to be associated with mating, feeding, aggression, and increased tolerance of social interactions. To test this, we estimated the abundances of neuropeptides of the burying beetles *N. vespilloides* sampled from solitary, active parenting, or a post-parenting and solitary state. Burying beetles, especially *N. vespilloides*, represent an excellent system to address the role of neuropeptides in parenting. Parenting is extensive and elaborate (Fig. 1). Adult beetles of this genus locate a vertebrate carcass and bury it. Parents then provide indirect care by removing the fur or feathers and forming a nest within the carcass. They also repeatedly coat the carcass with excretions that retard microbial growth. Direct parental care involves feeding larvae predigested carrion by regurgitation for the first two days of larval life (Fig. 1). Parenting occurs for 75% of larval development, yet lasts only days^19^ at which point larvae are fully-grown. *Nicrophorus vespilloides* is also molecularly tractable with a published genome^20^, allowing for efficient proteomic work and a characterization of the transcriptional response of a similar series of behavioural transitions. Finally, *N. vespilloides* is normally solitary but switches to parenting in the presence of suitable resources available (a vertebrate carcass) and restricted to a limited period of time. We can therefore sample females experimentally manipulated to be in non-overlapping behavioural states; from non-parenting and solitary, to parenting, or to post-parenting and solitary again^19^.

**Figure 1.**
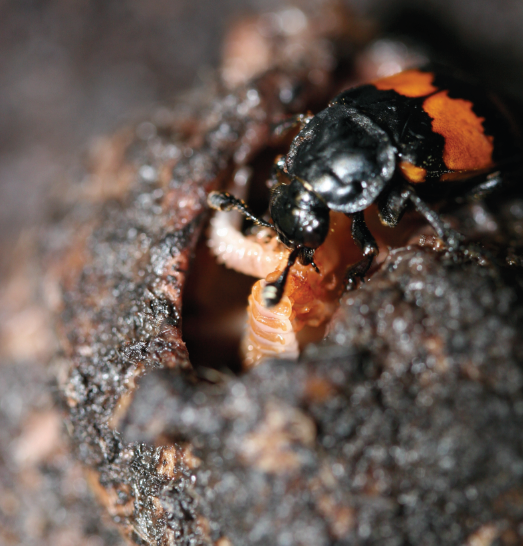
A female burying beetle feeding her begging, dependent offspring. In this species, a parent spends around 72 h preparing a carcass, after which larvae hatch and arrive at the carcass. Once larvae arrive, parents spend a further 72 h feeding larvae (with peak parenting 12-24 h after larval arrival), and then disperse around 100 h. Larvae disperse fully grown around 125 h after arrival on the carcass. As shown here, feeding involves direct mouth-to-mouth contact and a transfer of pre-digested carrion from the parent to the offspring. Photograph by A. J. Moore.

## Results

Our analysis identified 130 peptides in the brains of *N. vespilloides*. We found very few differences in the specific peptides that were identified for each neuropeptide proteins across the three behavioural states (i.e., peptides identified in one state but not others). Actively parenting individuals exclusively displayed two peptides from FMRFa: DKGHFLRF and GDLPANYEMEEGYDRPT. Actively parenting individuals exclusively displayed a single peptide from NPLP-1: KESYDDDYYRMAAF. No *Apis*-NVP-like peptides of the sequence FLNGPTRNNYYTLSELLGAAQQEQNVPLYQRYVL were found in actively parenting samples.

From these peptides we identified 17 neuropeptide proteins that were present in at least one behavioural state (Table 1, 2). Twelve were represented in all three behavioural states, while PBAN was absent in post-parenting individuals, ITP was restricted to virgins, sNPF was restricted to virgins and actively parenting, DH_47_ was restricted to actively parenting individuals, and CCAP was restricted to post-parenting individuals. Virgins showed a higher level of variability than the other two behavioural states.

**Table 1.**
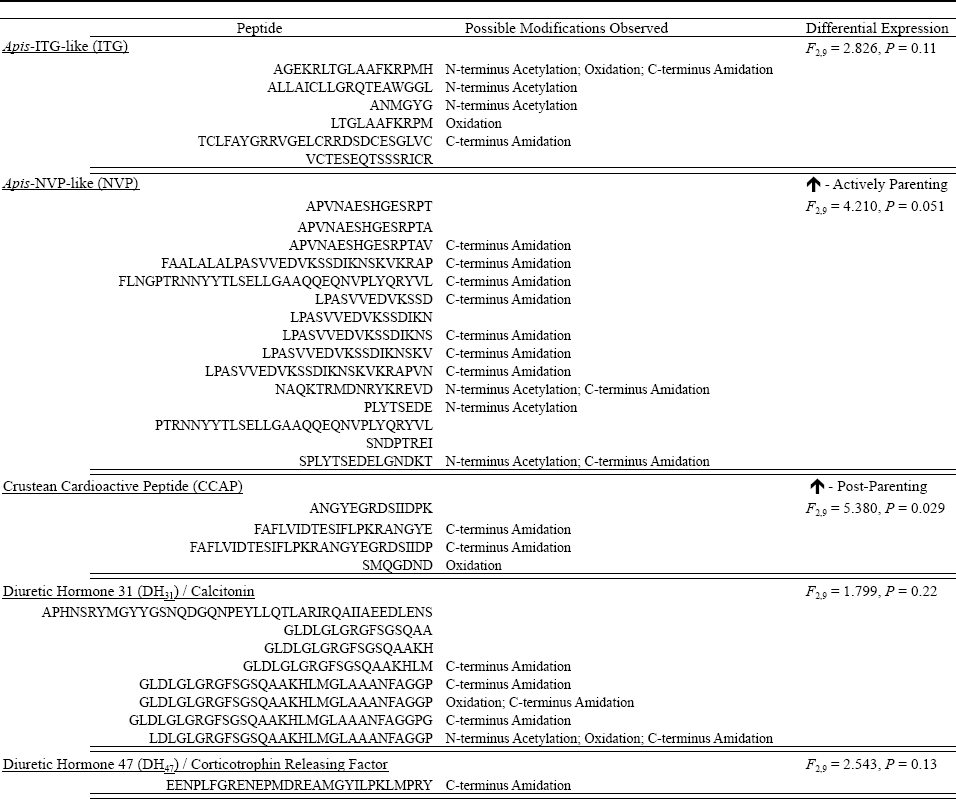
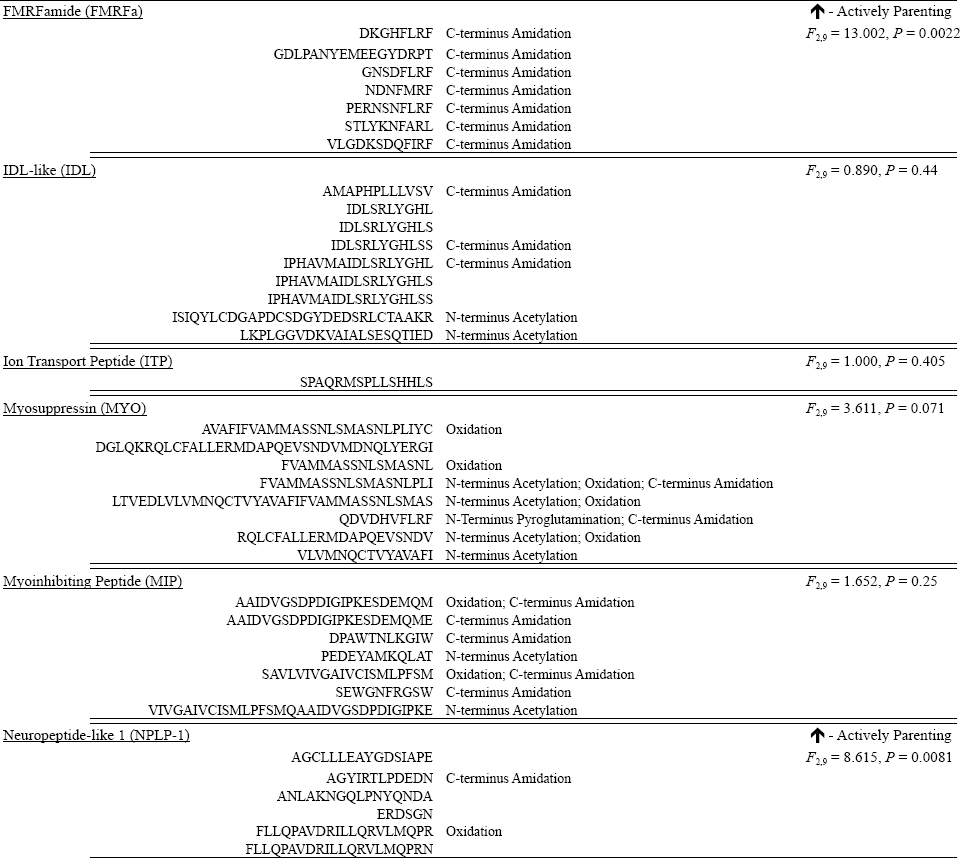
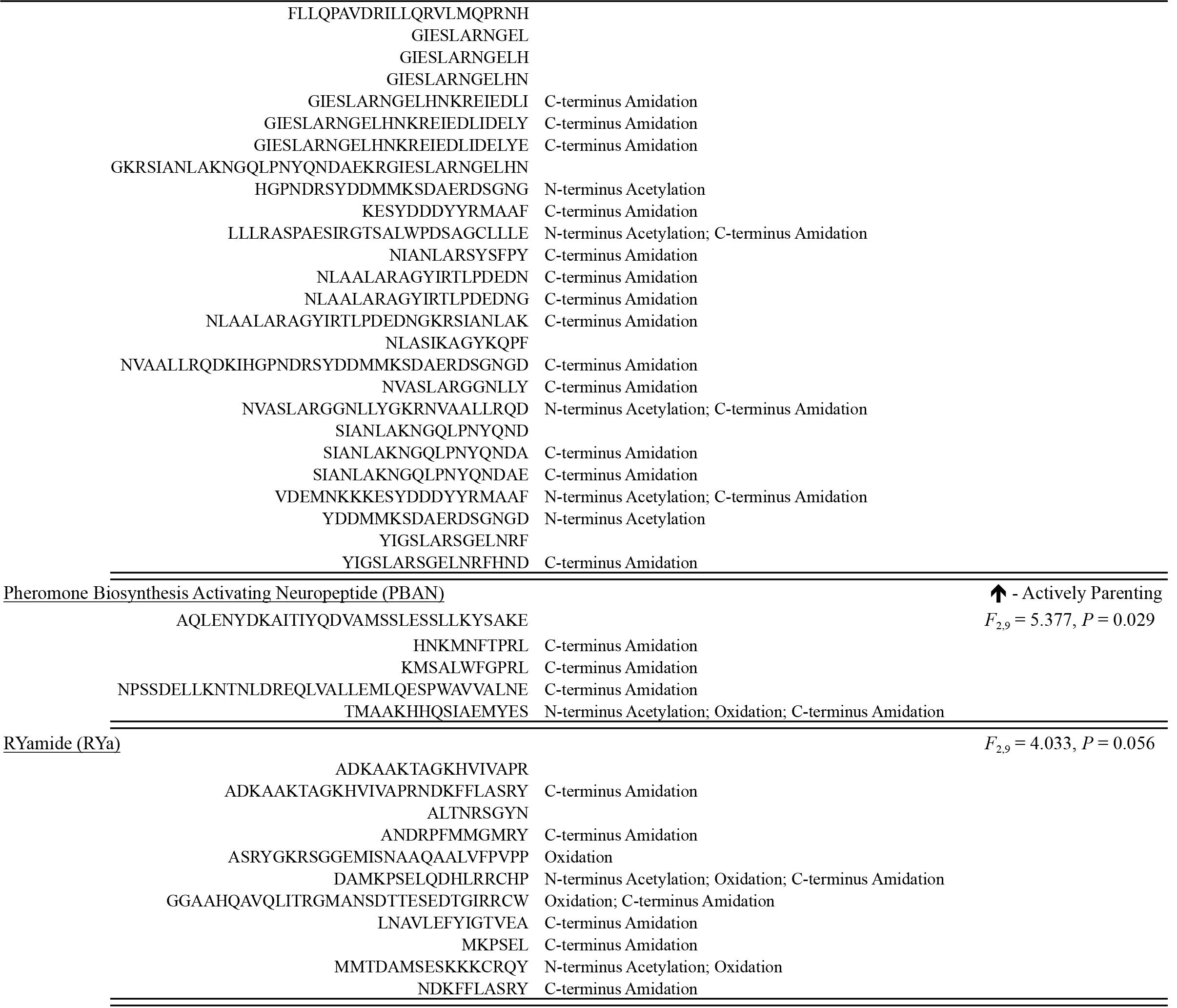
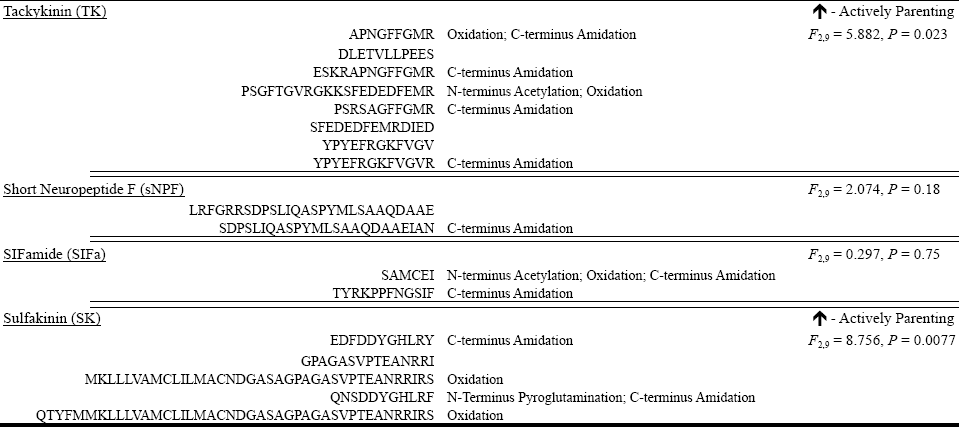
Individual peptides of the neuropeptide precursors identified with observed modifications and evidence of differential expression. Both functional and non-functional peptides are reported.

**Table 2.** Principal Component Analysis (PCA) of neuropeptide abundance of virgins, actively parenting and post-parenting *Nicrophorus vespilloides* females. PC’s with eigenvalues exceeding 1 are reported.

|  | Principal Component |  |  |  |  |
| --- | --- | --- | --- | --- | --- |
|  | PC1 | PC2 | PC3 | PC4 | PC5 |
| Neuropeptides |  |  |  |  |  |
| NPLP-1 | -0.325 | 0.184 | -0.001 | 0.052 | -0.009 |
| TK | -0.330 | 0.057 | -0.208 | -0.138 | 0.220 |
| *Apis* NVP-like | -0.271 | -0.174 | -0.208 | -0.230 | 0.149 |
| DH<sub>31</sub> | -0.185 | -0.086 | -0.287 | 0.442 | -0.376 |
| FMRFa | -0.284 | -0.186 | 0.261 | 0.292 | -0.011 |
| ITG | -0.246 | 0.268 | 0.148 | 0.164 | -0.363 |
| SIFa | -0.097 | 0.457 | -0.100 | -0.200 | -0.156 |
| IDL-like | -0.232 | 0.287 | -0.240 | -0.093 | -0.024 |
| SK | -0.281 | 0.144 | 0.385 | -0.029 | 0.181 |
| MYO | -0.261 | 0.216 | 0.362 | 0.150 | 0.246 |
| MIP | -0.177 | -0.348 | 0.151 | -0.356 | -0.156 |
| RYa | -0.186 | -0.368 | 0.298 | 0.224 | -0.034 |
| PBAN | -0.297 | -0.234 | -0.107 | 0.035 | 0.093 |
| ITP | 0.042 | -0.218 | 0.270 | -0.472 | -0.346 |
| sNPF | -0.209 | -0.181 | -0.306 | 0.045 | -0.419 |
| DH<sub>47</sub> | -0.211 | -0.225 | -0.315 | -0.082 | 0.418 |
| CCAP | 0.286 | -0.136 | -0.079 | 0.369 | 0.184 |
| Eigenvalues | 6.679 | 3.497 | 1.896 | 1.612 | 1.401 |
| % Variance Explained | 39.28 | 20.57 | 11.15 | 9.48 | 8.24 |

Having defined these neuropeptides, we tested for changes in the relative abundances of all neuropeptides across the three behavioural states tested (virgins, actively parenting, and post-parenting individuals) using a multivariate analysis of variance (MANOVA). We found statistically significant overall differences in the relative abundance between the states (*F*_2,9_ = 27.678; *P* = 0.0001) The main difference reflects level of expression in the different states (Fig. 2, Table 2). We next used univariate comparisons (ANOVA’s) to examine how the relative abundances of specific neuropeptides were changed. We found six neuropeptides were more highly expressed with actively parenting individuals. NPLP-1 was differentially expressed (*F*_2,9_ = 8.615, *P* = 0.0081), with statistically significantly higher expression of actively parenting compared with post-parenting (*P* = 0.0063). TK was differentially expressed (*F*_2,9_ = 5.882, *P* = 0.023), also with statistically significantly higher expression when individuals were actively parenting compared with post-parenting (*P* = 0.020). FMRFa was differentially expressed (*F*_2,9_ = 13.002, *P* = 0.0022), also with statistically significantly higher expression when individuals were actively parenting compared with virgins (*P* = 0.011) and post-parenting (*P* = 0.0023). SK was differentially expressed (*F*_2,9_ = 8.756, *P* = 0.0077), with statistically significantly higher expression in virgins (*P* = 0.026) and actively parenting (*P* = 0.0087) compared with post-parenting. PBAN was differentially expressed (*F*_2,9_ = 5.377, *P* = 0.029), with statistically significantly higher expression when individuals were actively parenting compared with post-parenting (*P* = 0.023). NVP was differentially expressed (*F*_2,9_ = 4.210, *P* = 0.051), with higher expression when individuals were actively parenting compared with post-parenting (*P* = 0.043). One neuropeptide, CCAP had statistically significantly lower expression in parenting individuals (*F*_2,9_ = 5.380, *P* = 0.029), with higher expression in post-parenting than in either virgins (*P* = 0.046) or actively parenting (*P* = 0.046).

**Figure 2.**
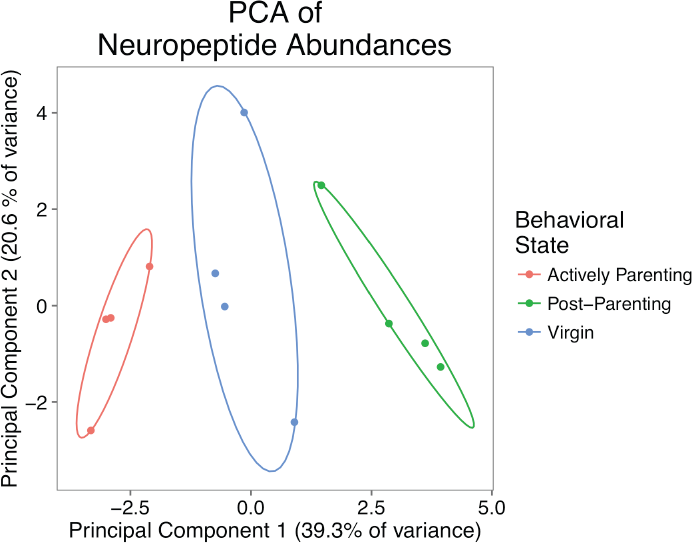
Principal component analysis (PCA) of all neuropeptide relative abundances. Graph of the association between abundances and three non-parenting and parenting behavioural states of *Nicrophorus vespilloides.* Ellipses show the 95% confidence area of each group.

While not reaching the level of conventional statistical significance, we identified two neuropeptides that showed a strong trend toward differential expression. RYa (*F*_2,9_ = 4.033, *P* = 0.056) and MYO (*F*_2,9_ = 3.611, *P* = 0.071) were also most highly expressed in actively parenting individuals. The remaining neuropeptides showed no strong trends. There was no suggestion of differential expression of DH_31_ (*F*_2,9_ = 1.799, *P* = 0.22), ITG (*F*_2,9_ = 2.826, *P* = 0.11), SIFa (*F*_2,9_ = 0.297, *P* = 0.75), IDL (*F*_2,9_ = 0.890, *P* = 0.44), MIP (*F*_2,9_ = 1.652, *P* = 0.25), ITP (*F*_2,9_ = 1.000, *P* = 0.405), sNPF (*F*_2,9_ = 2.074, *P* = 0.18), and DH47 (*F*_2,9_ = 2.543, *P* = 0.13).

## Discussion

Our goal was to test the prediction that the mechanisms involved in the evolution of parental care reside in predictable pathways reflecting co-opted behavioural precursors^1^,^2^. To do this we examined peptide abundance, with the prediction that the neuropeptides differentially expressed during parenting would function in feeding, mating, aggression, and social interactions in organisms that do not display parental care. We profiled these changes from brains of the burying beetle *Nicrophorus vespilloides*, which provides direct care by regurgitating food to dependent offspring. We identified 17 neuropeptides in the brain of *N. vespilloides*, which is consistent with other studies of non-model organisms^21-23^. Of these, seven were differentially expressed, with six up-regulated during parenting, in our comparison of the neuropeptides of individuals not parenting or post-parenting.

Parenting across species typically involves a pause of mating, feeding others, appropriately directed aggression for defence, and social interactions^1-3^. The six neuropeptides that were differentially expressed (Table 1) support this prediction of these co-opted pathways. In other insects, both FMRFa and SK influence mating^24,25^. Feeding behaviour and food intake are influenced by NVP and SK^22,26-28^. Aggression and resource defence are influenced by TK^29,30^ and SK^25^. NPLP-1, TK, and PBAN all influence tolerance of social interactions^21,31,32^. Of the 11 neuropeptides that were not differentially expressed, many have poorly understood functions (e.g., ITG, RYa, MIP, MYO^25,33,34^), or function outside the predicted pathways (CCAP, DH_31_, DH_47_, IDL, ITP^34^). Two of these neuropeptides have the potential to function in the predicted pathways were sNPF, which influences feeding, and SIFa, which influences reproduction^25,34,35^. Critically, none of the differentially expressed neuropeptides we identified in this study function solely outside the predicted pathways. Thus, like candidate gene studies^11^, hypotheses about pathways are likely to be more robust than hypotheses focused on specific neuropeptides when examining homologous behaviour in novel species.

Our study suggests three areas for further consideration to understand the mechanisms underlying parental care. First, we suggest that knowing the selective pressures leading to behavioural evolution provides insights into mechanisms by providing predicted pathways is general. This can be tested in other behaviours where the selective pressures are known and therefore the underlying behavioural traits that are predicted to change can be identified *a priori*. Second, we provide information about specific neuropeptides that appear to underpin parental care and these can be examined in other subsocial organisms. Functional studies are desperately needed for organisms outside the genetic model species. Finally, by specifying the behavioural and genetic pathways expected to be co-opted when parenting evolves, we can then identify particularly influential molecules that deserve further examination in *N. vespilloides*. Among those neuropeptides we have identified, both tachykinin and sulfakinin influence nearly all of the pathways thought to be co-opted during the evolution of parenting and deserve further investigation.

## Methods

### Experimental Design

We used female *N. vespilloides* derived from an outbred colony we maintain at the University of Georgia, Athens. The colony was founded with beetles originally captured from Cornwall, UK and is subsidized yearly with new beetles from the same location. Beetles were fed once weekly with decapitated mealworms *ab libitum* and kept on a 15:9 hour light:dark cycle. Further details of colony maintenance can be found in Cunningham et al.^36^ with the exception of a change of soil type (to Happy Frog potting soil, FoxFarm, Arcata, CA, USA).

To examine how neuropeptide expression changed with transitions of behavioural state, we collected age-matched females in three behavioural states: virgin (no social experience, no mating, no reproductive resource, and no parenting), actively parenting (social experience, mated, reproductive resource, and actively parenting), post-parenting (social experience, mated, reproductive resource, and past parenting experience). Full descriptions of each behavioural state can be found in Roy-Zokan et al.^37^ We collected virgins directly from their individual housing boxes. We collected actively parenting females directly from the carcass cavity where offspring are fed. We collected post-parenting females nine days from the start of a breeding cycle after they had been isolated for 24 hours. We collected all beetles at 19-22 days post-adult eclosion and all beetles were fed one day before their collection or before their pairing to standardize feeding status.

We performed dissections in ice-cold 1x PBS (National Diagnostics, Atlanta, GA, USA) and completed them within four minutes. We placed single brains into 0.6 mL Eppendorf tubes with 30 μL of ice-cold acidified acetone extraction buffer (40:6:1 (v/v/v) Acetone: H_2_O: Concentrated HCl). We did not collect the retro-cerebral complex (corpora allata-corpora cardiaca). Once collected, we stored samples at -80 °C until extraction.

We pooled eight brains into a single biological replicates by removing brains and their associated acetone extraction buffer to a single 2.0 mL low protein binding Sartorius Vivacon 500 tubes (Göttingen, Germany). We collected four biological replicates per behavioural state. We sonicated each biological replicate with a Misonix Sonicator S-4000 (Farmingdale, NY, USA) fitted with a 1/8’’ tip (#419) set to an amplitude of 20 for a total of 60s sonication with 15s pulses followed by 15s rest on ice. We then centrifuged replicates at 16,000 g for 20 minutes at 4 °C with a 5810-R Eppendorf centrifuge. We collected the supernatant into a new Vivacon tube and repeated the extraction with the same volume of buffer and sonication protocol. We pooled and extracted all replicates at the same time without ordering. We stored samples at 4 °C until LC-MS/MS analysis.

We analysed our biological replicates with a Finnigan LTQ linear ion trap mass spectrometer (Thermo-Fisher) and an 1100 Series Capillary LC system (Agilent Technologies) with an ESI source with spray tips built in-house. The extraction buffer was vacuum-dried off of all biological replicates with a VirTis Benchtop K Lyophilizer (SP Scientific, Warminster, PA, USA) and biological replicates were suspended in 11 μL of buffer A [5% acetonitrile/0.1% formic acid/10 mM ammonium formate] and 8 μl of each replicate were injected into the LC column. Peptides were separated using a 200-μm x 150-mm HALO Peptide ES-C18 column packed with 5-μm diameter superficially porous particles (Advanced Materials Technology). The gradient used for each replicate was 5-75% buffer B (80% acetonitrile/0.1% formic acid/10 mM ammonium formate) for 120 minutes at a 2 μL/min flow rate. The settings for the mass spectrometer included taking the 5 most intense ions from each full mass spectrum for fragmentation using collision-induced dissociation (CID) and the resulting MS/MS spectra were recorded. Our biological replicates from the three treatments were interspersed with each other for LC-MS/MS analysis. All chemicals were LC-MS or molecular biology grade.

### Neuropeptide Identification and Analysis

We converted the resulting RAW spectra using Trans Proteomic Pipeline (Seattle Proteome Center, Seattle, WA, USA). MS/MS spectra were then imported into MASCOT (v2.2.2; MatrixScience, Boston, MA, USA) and searched against all annotated proteins from the *N. vespilloides* genome^20^. We set search parameters as: enzyme, none; fixed modifications, none; variable modifications as oxidation (M), acetyl (N-terminus), pyroglutamic acid (N-terminus Glutamine), and amidation (C-terminus); maximum post-translational modifications, 6; peptide mass tolerance, ± 1000 ppm; fragment mass tolerance, ± 0.6 Da.

We imported MASCOT results into ProteoIQ (v2.6.03; Premier Biosoft, Palo Alto, CA, USA) to estimate abundance of neuropeptides. We identified proteins, peptides, and assigned spectral counts using all biological replicates for each behavioural state. This analysis produces a list of peptides assigned to each identified protein and from this we looked for qualitative differences in the presence/absence of peptides across the behavioural states for peptides that had at least three spectra and were not truncated forms of a larger observed peptide from a particular protein. We excluded peptides from proteins that were only observed in a single behavioural state. We then calculated normalized spectral abundance factor (NASF’s) for all proteins within each biological replicate using the protein length for the NASF length correction factor^38^. Only peptides with at least two spectra within one biological replicate were quantified. Neuropeptide proteins were extracted from the overall protein list after establishing their identity within the published *N. vespilloides* gene set with a *Tribolium castaneum* neuropeptidome^39^ and confirming their identity using NCBI’s non-redundant insect protein database.

To test the hypothesis that changes in neuropeptide expression can be predicted *a priori*, we first performed a MANOVA to establish that there was an overall difference in the neuropeptide composition between treatments. We followed this multivariate test with univariate tests (ANOVAs) for difference of individual neuropeptide abundance, testing for the effect of behavioural state on expression. We performed *post-hoc* tests of differences in the pairwise means of the behavioural states using Tukey-Kramer HSD tests. All statistical analyses were conducted with JMP Pro (v11.0.0, Cary, NC, USA). Visualizations were prepared in R (v3.2.1) using prcomp function and ggbiplot (github.com/vqv/ggbiplot).

## Acknowledgements.

We thank Elizabeth McKinney for technical support. This research was supported by funding to AJM from the US National Science Foundation (IOS-1354358) and the University of Georgia’s Office of the Vice-President for Research.

## Author Contributions.

CBC, RBM, RO, AJM designed the experiment. CBC gathered and extracted the biological samples with assistance from RBM. MJB and RO conducted the LC-MS/MS analysis. CBC and AJM analysed results with guidance from MJB and RO. CBC and AJM wrote the manuscript with input from all authors. All authors approved the final manuscript.

## Conflict of Interest

We declare no conflict of interest.

